# Intestinal epithelial Axin1 deficiency protects against colitis via altered gut microbiota

**DOI:** 10.1101/2022.03.23.485334

**Authors:** Shari Garrett, Yongguo Zhang, Yinglin Xia, Jun Sun

**Author notes:** **Corresponding author** Jun Sun, Ph.D., AGAF, FAPS, Professor. **Author Contributions** SG performed the cellular and animal studies, the detailed analyses of the results; YZ: animal studies; SG and JS, prepared the figures and the draft text; YX contributed to the statistical analysis of data and the draft text; and JS obtained funds, designed the study, and directed the project. All authors contributed to the writing of the manuscript.

## Abstract

**Background and Aims:** Intestinal homeostasis is maintained by specialized host cells and the gut microbiota. Wnt/β-catenin signaling is essential for gastrointestinal development and homeostasis, and its dysregulation has been implicated in inflammation and colorectal cancer. Axin1 negatively regulates activated Wnt/β-catenin signaling, but little is known regarding its role in regulating host-microbial interactions in health and disease. Here, we aim to demonstrate that intestinal Axin1 determines gut homeostasis and host response to inflammation.

**Methods:** The expression of Axin1 was analyzed in human IBD datasets. To explore the effects and mechanism of intestinal Axin1 in regulating intestinal homeostasis and colitis, we generated mouse models with Axin1 conditional knockout in intestinal epithelial (Axin1^ΔIEC^) and Paneth cells (Axin1^ΔPC^) to compare with control (Axin1^LoxP^) mice.

**Results:** We found increased Axin1 expression in the colonic epithelium of human IBD. Axin1^ΔIEC^ mice exhibited altered goblet cell spatial distribution, Paneth cell morphology, reduced lysozyme expression, and enriched *Akkermansia muciniphila*. Absence of intestinal epithelial and Paneth cell Axin1 decreased susceptibility to DSS-induced colitis *in vivo*. Axin1^ΔIEC^ and Axin1^ΔPC^ became more susceptible to DSS-colitis after cohousing with control mice, suggesting the non-colitogenic effect is driven by the gut microbiota.

**Conclusion:** We found loss of intestinal Axin1 protects against colitis, which is likely driven through Paneth cell Axin1 and the microbiota. Our study demonstrates a novel role of Axin1 in mediating intestinal homeostasis and the microbiota. Further mechanistic studies using specific Axin1 mutations elucidating how Axin1 modulates microbiome and host inflammatory response, will provide new therapeutic strategies for human IBD.

**What you Need to Know:** *Background and Context:* Wnt/beta-catenin is a fundamental molecular pathway that affects intestinal proliferation and differentiation. Axin1 negatively regulates activated Wnt/β-catenin signaling, but little is known regarding its role in the microbiome. Dysfunction of Wnt/beta-catenin was reported in human inflammatory bowel disease (IBD) and Axin1 serum level was elevated in patients with UC.

*New Findings:* We found increased Axin1 expression at both the mRNA and protein level in human IBD. Specifically, we identified increased Axin1 expression positive correlated with pro-inflammatory cytokines IL-6 and TNF-α in CD. Our study, for the first time, identifies links between the gut microbiota and intestinal Axin1 in intestinal inflammation through utilization of innovative deletion mouse models in intestinal epithelium and Paneth cells. Loss of intestinal Axin1 plays a novel role in intestinal inflammation by altering the Paneth cells and microbiome (e.g., enriched *Akkermansia mucinlphila)*. Our study has provided insights into the molecular mechanism that might contribute to IBD, especially the novel role of Paneth cell Axin1 in colitis.

*Limitations:* There are no human or mice studies assessing the role of intestinal epithelial and Paneth cell Axin1 in inflammation and the microbiome.

*Impact:* Further explorations of the gut microbiota and Axin1 interaction as we report will provide novel mechanistic strategies for therapeutic approaches for human IBD by targeting intestinal Axin1 and Axin1-associated microbiome.

## Introduction

IBD is a chronic inflammation of the intestines affecting more than 6 million people globally [1]. The pathogenesis of IBD remains unclear but has been observed to involve complex interactions of environmental factors, gut microbiota, host epithelial cells and dysregulation of innate and adaptive immunity against the backdrop of genetic pre-disposition [2]. A major pathway implicated in IBD, is the dysregulation of intestinal Wnt/β-catenin signaling [3, 4, 5, 6]. Wnt/β-catenin signaling is an essential regulator of gastrointestinal development and homeostasis [7]. Axin1 was initially identified as an inhibitor of Wnt/β-catenin signaling and a regulator of embryonic axis formation [8]. Axin1 is a scaffold protein that recruits the β-catenin destruction complex, facilitating GSK3β’s phosphorylation of β-catenin, targeting it for ubiquitination and degradation by the proteasome [9, 10, 11]. However, the role of Axin1 in regulating intestinal inflammation and the development of IBD is unknown.

Axin1 has a distinct biological role in bacterial infections, specifically host-pathogen interactions [12]. *Salmonella* decreases Axin1 protein expression in intestinal epithelial cell lines at the post-transcriptional level, whereas its overexpression inhibits *Salmonella* invasion and inflammation *in vitro* [12]. However, the role of intestinal Axin1 signaling in maintaining mucosal health has not been fully understood *in vivo*. Since whole body Axin1 deletion is lethal, few investigators are utilizing conditional Axin1 knockout mouse models for mechanistic studies [8].

In the current study, we hypothesize that intestinal epithelial Axin1 plays a role in regulating the microbiota and susceptibility to inflammation. In human IBD samples, we found increased expression of Axin1 at the mRNA and protein levels. To investigate the molecular mechanism of intestinal epithelial Axin1 regulation, we generated a novel mouse model of intestinal epithelial cell conditional knockout of Axin1 (Axin1^ΔIEC^) [13]. Altered expression of intestinal Axin1 impairs intestinal epithelial secretory cells and cell differentiation. Furthermore, we generated a mouse model of Axin1^ΔPC^ to study the tissue-specific role of Paneth cell Axin1 in response to inflammation. Our studies for the first time demonstrate the relationship between intestinal Axin1 and the maintenance of intestinal and microbial homeostasis. Understanding complex interactions among host factors (e.g., Axin1), cellular changes, and the microbiota in colitis will help provide novel therapeutic approaches for human IBD.

### Material and Methods

#### Human Intestinal Biopsies

Slides containing paraffin-embedded colon biopsy samples of patients with UC, CD and healthy controls were obtained from a tissue microarray from US Biomax, Inc. (CO246).

#### Gene Expression Datasets

We used microarray data registered on Gene Expression Omnibus repository (https://www.ncbi.nlm.nih.gov/geo/). From the GEO repository, colonic mucosal biopsies from inflamed mucosa of IBD patients with ulcerative colitis, Crohn’s colitis, and healthy controls. Total RNA was isolated via microarray and reported (GEO accession number GSE 16879). Gene expression data for Axin1 from UC controls (n = 5), UC (n = 22), CD controls (n = 5) and CD (n = 12) patients were extracted and analyzed for our study.

#### Experimental Animals

Axin1^loxp/loxp^ (Axin1^LoxP^) mice were originally reported by Xia et al. [8]. Axin1^ΔIEC^ mice were obtained by crossing Axin1^LoxP^ with Vil1/villin-Cre mice (Jackson Laboratory, 004586, Bar Harbor, ME, USA). Defa6-cre mice were from Dr. Blumberg [14]. Axin1^ΔPC^ mice were obtained by crossing Axin1^LoxP^ mice with DEFA6-cre mice. Experiments were performed on 6–8-week-old mice and were provided water ad libitum and maintained in a 12 h dark/light cycle. All animals were housed in the Biologic Resources Laboratory (BRL) at the University of Illinois at Chicago (UIC) and utilized in accordance with UIC Animal Care Committee (ACC) and Office of Animal Care and Institutional Biosafety (OACIB) guidelines. Animal work was approved by the UIC Office of Animal Care (ACC15-231, ACC17-218, and ACC18-216).

#### Genetic Background of Mouse Strains

Axin1 targeting vector was electroporated into SV129 embryonic stem cells. These embryonic stem (ES) cells were selected and assessed for Axin1 disruption at exon 2. The proper ES clones were injected into C57BL/6J blastocysts to produce chimeric animals. Chimeric animals were further crossed to produce Axin1^loxp/loxp^ (Axin1^LoxP^) mice which were viable and fertile with no recognizable phenotype. Villin-cre (stock no: 004586) transgenic mice were purchased from Jackson Laboratory (Bar Harbor, ME, USA), and had been generated on a C57BL/6 background. The Paneth cell specific defa6-cre mouse strain [14] was generated by injecting the designed plasmid into the pronucleus of C57BL/6 mice, allowing for cre-recombinase to be expressed under Paneth cell specific promotor, defensin-6.

#### Colitis Induction

Colitis was induced as previously described [15, 16]. Mice were administered 5% DSS (MW = 40-50 kDa; USB Corp. Cleveland, OH) dissolved in filter purified water ab libitum during the experimental period. Animals were weighed daily. At day 7 mice were sacrificed under anesthesia and the severity of colitis was quantified by a disease activity index, determined by percent of weight loss, fecal blood, and diarrhea.

#### Co-housing Experiment

Male and female Axin1^LoxP^ and Axin1^ΔIEC^ or Axin1^ΔPC^ mice (6-to 8-week old) were co-housed in new cages. Each cage contained 3 Axin1^LoxP^ mice and 2 Axin1^ΔIEC^ or 2 Axin1^ΔPC^ mice as previously described [17]. After 4 weeks of co-housing, 5% DSS dissolved in filter-purified drinking water was given ab libitum. Animals were weighed daily. At day 7 after DSS administration mice were sacrificed under anesthesia and the severity of colitis was quantified by a disease activity index, determined by percent of weight loss, fecal blood, and diarrhea.

#### Histology of mouse colon and small intestine

Intestines were harvest as previously described [15, 16, 18, 19] and fixed in 10% formalin (pH 7.4), processed and paraffin embedded. 4µm sections were stained with hematoxylin and eosin [20]. Histological damage was scored as described [13]. Alcian Blue staining was performed according to standard protocol using alcian blue pH 2.5.

#### Immunoblotting

Mouse ileal and colonic epithelial cells were collected by scraping the tissue and homogenized as previously described [17, 21]. Equal amounts of normalized protein were separated by SDS-polyacrylamide gel electrophoresis, transferred to nitrocellulose, and immunoblotted with primary antibodies as previously described [12, 22]. Antibodies were visualized by ECL. Membranes probed with more than one antibody were stripped before re-probing.

#### Immunohistochemistry

Intestinal tissues were fixed in 10% buffered formalin and processed with standard techniques as previously described [15, 16, 22, 23]. Slides were stained with anti-Axin1(Invitrogen, 34-5900) and staining intensity was performed as previously described [24].

#### Immunofluorescence

Intestinal tissues were freshly isolated and embedded in paraffin wax after fixation with 10% neutral buffered formalin. Immunofluorescence was performed on paraffin-embedded sections (4 μm) of mouse intestine. After preparation of slides as described previously [15], sections were incubated with anti-lysozyme (Santa Cruz Biotechnology Inc., CA) antibody overnight at 4°C. Samples were then incubated with donkey-anti goat Alexa Flour 488 (Thermo Fisher Scientific, D1306) for 1 hour at room temperature. Tissues were mounted with SlowFade (Thermo Fisher Scientific, S2828), cover slipped and were sealed. Sections were examined with a Leica SP5 Laser Scanning confocal microscope (LSM 710, Carl Zeiss Inc., Germany) or Olympus BX51 Fluorescence Microscope (Olympus Life-Sciences, PA).

#### Fluorescence in situ Hybridization

Fluorescent in situ hybridization (FISH) was performed using anti-sense ssDNA probe EUB 388 (5′-GCT GCC TCC CGT AGG AGT-3′) targeting a highly conserved region of the bacterial 16s gene. 4 μm tissue sections were baked for 30 minutes at 60°C. Sections were deparaffinized in xylene, dehydrated with 100% ethanol, dried and incubated in 0.2M HCl for 20 minutes and heated in 1M NaSCN for 10 minutes at 80°C. Sections were digested with pepsin (4% pepsin in 0.01M HCl) for 20 minutes at 37°C. Slides were washed in wash buffer (0.3M NaCl, 0.03 M sodium citrate, pH 7.0). Sections were fixed for 15 minutes in 10% buffered formalin. Probes were hybridized at 5ng/µl for 5 minutes at 90°C in hybridization buffer (0.9M NaCl, 30% formamide, 20 mMTris-HCl (pH 7.4) and 0.01% SDS and incubated overnight at 37°C. Slides were washed 5 times for 5 minutes at 45°C in wash buffer. To visualize cell nuclei sections were stain with DAPI/antifade solution. Slides were examined with Olympus BX51 Fluorescence Microscope (Olympus Life-Sciences, PA).

#### Transmission Electron Microscopy

Small intestines were fixed in 4% paraformaldehyde/3% glutaraldehyde, in 10mM sodium phosphate buffer (pH 7.4) 48 hours. All samples were prepared as previously described [15]. After the resin was polymerized, samples were sliced into 1×2×2 mm pieces and examined with a Philips CM 100 electron microscope (Philips, Netherlands) at an accelerating voltage of 80 KV for imaging at the UIC electron microscopy core.

#### Multiplex ELISA Assay

A mouse specific ProcartalPlex Multiplex Immunoassay Plate (Invitrogen Thermo Fisher) was used to detect serum cytokine levels. The assay was performed using the manufacturer’s instruction manual using proper standards. Plates were read using a Magpix Luminex Machine.

#### Isolation of Paneth cells

Small intestines were harvested. Paneth cells were isolated as previously described [15]. After centrifugation, cells were resuspended in 2mM EDTA-1% FBS and incubated with CD24-PE antibody in the dark at 4°C for 30 minutes on a shaker. Cells were centrifuged and washed 2 times in flow wash buffer and then resuspended in flow wash buffer for sorting.

#### Bioinformatic Analysis of 16S rRNA Sequencing of Fecal Data

Fecal samples were harvested and prepped as previously described [17]. Samples were analyzed at the UIC Genome Research Core utilizing the Illumina sequencing platform. The QIIME pipeline[25] was used to process raw sequence data, including read merging, adapter &quality trimming, and chimeric checking and to generate operational taxonomic units (OTUs) in closed reference manner using the UCLUST method with a threshold of 97% sequence similarity. Taxonomic annotations were assigned to each OTU using the representative sequence data against the Illumina Curated GreenGenes reference database [26].

#### Real-Time Quantitative PCR

Total RNA was extracted from mouse colonic and small intestinal epithelial cells using TRIzol reagent. RNA reverse transcription was done using iScript cDNA synthesis kit according to the manufacturer’s directions (Bio-Rad, Heracules, CA). cDNA reaction products were subjected to quantitative real-time PCR using the iQ SYBR green supermix according to the manufacturer’s instructions. All expression levels were normalized to villin levels in the same sample. Percent expression was calculated as the ratio of the normalized value of each sample to the corresponding control. All real-time PCR reactions were performed in duplicate.

#### Real-Time Quantitative PCR of Bacterial DNA

DNA was extracted from mice feces using the stool DNA Kit (Omega Bio-tek, Norcross, GA) according to the manufacturer’s instructions. 16S ribosomal DNA PCR reactions were performed on the CFX Connect Real Time System (Bio-Rad Laboratories) and amplified using iTaq Universal SYBR green supermix (1725121; Bio-Rad Laboratories) according to the manufacturer’s directions. Primers specific to 16S rRNA were used as endogenous control to normalize between samples. The relative amount of 16S ribosomal DNA in each sample was estimated using the ΔΔCT method. Primer sequences were designed using Primer-BLAST or obtained from Primer Bank pairs listed.

#### Statistical Analyses

Data is expressed as mean ± SEM with p values ≤ 0.05 were considered as statistically significant. Shapiro-Wilks normality test was performed to detect whether the data significantly departures from normality. Parametric or non-parametric analyses were determined based whether the variables are normally distributed or not. An *F*-test of equality of variances was performed to test the null hypothesis that two normal populations of groups have the same variance. Differences between two groups with normal distribution were analyzed by unpaired student’s *t*-test for equal variances and Welch’s *t*-test for unequal variances respectively. Differences between two groups with non-normal distribution were analyzed by Wilcoxon rank sum test. Differences among three or more groups were analyzed by one-way ANOVA or two-way ANOVA as appropriate. To adjust for multiple comparisons, p-values were adjusted by the Tukey’s method. Spearman correlation analysis and scatter plots were performed to detect the correlation between Axin1 and IL-6, and TNF-*α* cytokines. Taxonomic and OTU abundances were analyzed as done in previous book [27]. Briefly, Principal Coordinate Analysis (PCA) and Shannon diversity analysis were conducted using R packages phyloseq and vegan. Statistical analyses were performed using GraphPad Prism version 8.0.0 for Windows (GraphPad Software, San Diego, California USA) and R software (R Core Team (2021), R Foundation for Statistical Computing, Vienna, Austria).

## Results

### Increased Axin1 Expression in Human IBD

The status of Axin1 in the inflamed intestine is unexplored. We analyzed microarray data of human colonic samples using the Gene Expression Omnibus database. We found the mRNA expression levels of Axin1 were elevated in human ulcerative colitis (UC) and Crohn’s disease (CD) (Fig 1A). Spearman correlation analysis indicated a positive correlation of Axin1 expression and pro-inflammatory cytokines IL-6 (Fig 1B) and TNF-α (Fig 1C) in CD. To investigate the changes and localization of Axin1 at the protein level, we performed immunohistochemistry of colonic tissue from healthy control, UC, and CD subjects. UC (Fig 1D) and CD (Fig 1E) colonic tissue showed a significant increase in Axin1 expression in the inflamed mucosa compared to normal colon.

**Figure 1:**
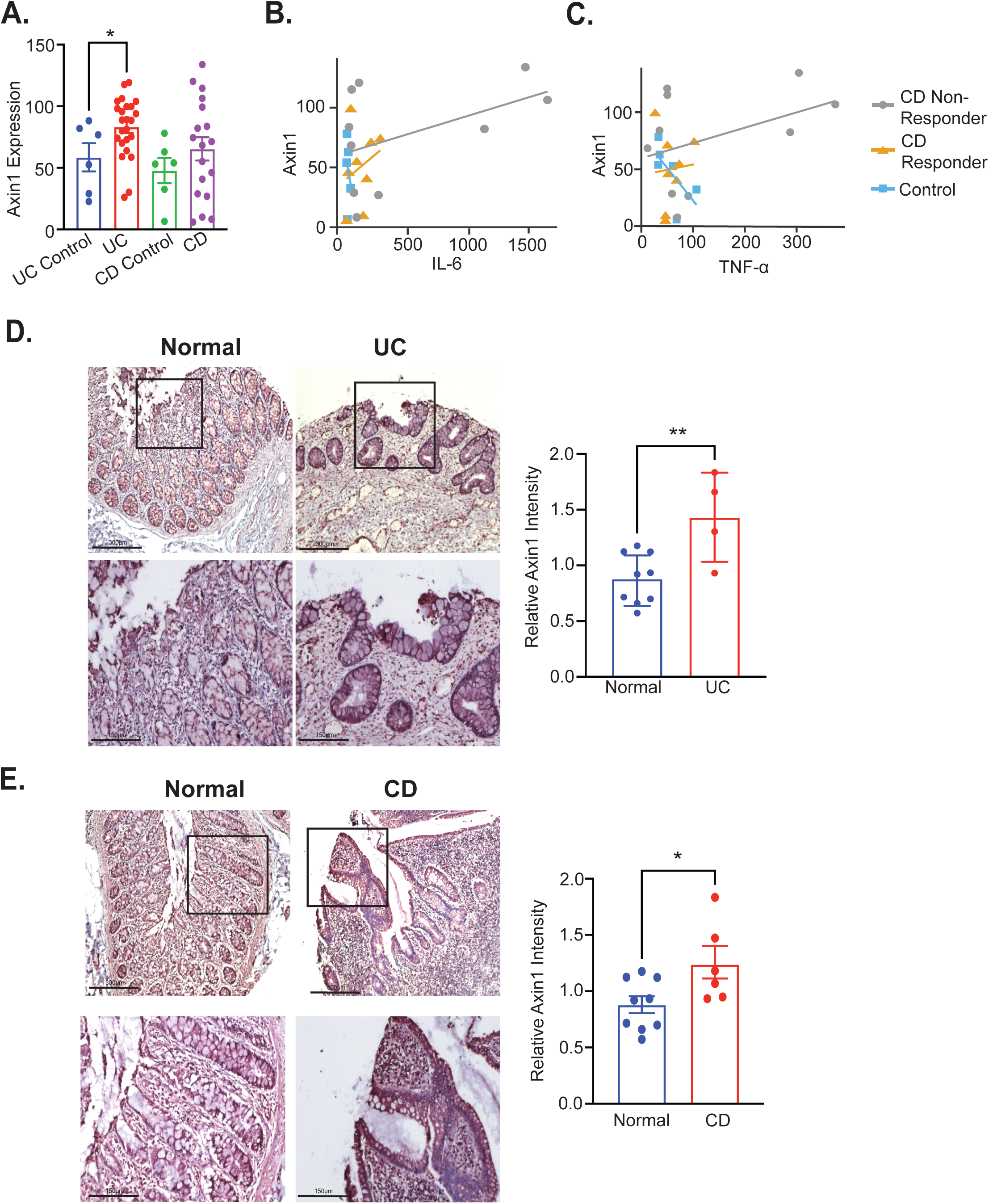
Expression of Axin1 is upregulated in human IBD. Axin1 mRNA expression in patients with UC and CD. Values for healthy control, CD and UC patients were obtained from Gene Expression Omnibus database GSE 16879. Data is expressed as mean+/-SEM; controls n = 5; UC, n = 22; CD = 16; One Way-ANOVA, *P<0.05. (B) Significantly coordinated expression of Axin1 and IL-6 or (C)TNF-α in CD patients. We performed a correlation analysis of Axin1 against IL-6 or TNF-α in GEO database GSE 16879 of patients who responded (CD Responder) and did not respond (CD Non-Responder) to the IBD drug infliximab prior to treatment, control n= 6; CD Responder n = 8; CD Non-Responder n =10, p = 0.0157 (IL-6) and p = 0.0563 (TNF-α). (D) Immunohistochemistry staining of Axin1 protein in colon of UC patients (n = 4) and healthy controls (n =9). (E) Immunohistochemistry staining of Axin1 protein in colon of CD patients (n = 6) and healthy controls (n =9). Student’s unpaired t-test, *P<0.05, and **P<0.01.

### Establishment of an Axin1^ΔIEC^ Mouse Model

We hypothesized that intestinal epithelial Axin1 plays a role in the pathogenesis of colitis. We generated a novel conditional IEC knockout by crossing an Axin1^LoxP^ mouse strain with villin-cre mice (Fig 2A). We checked expression of Axin1 mRNA in the mouse colon and small intestine. There was a significant decrease of Axin1 mRNA in the Axin1^ΔIEC^ mice (Fig 2B). We analyzed the protein level via western blot using intestinal mucosal scrapings. We found a significant reduction of intestinal epithelial Axin1 in both small intestine and colon of Axin1^ΔIEC^ mice (Fig 2C). We examined the localization of Axin1 expression in the intestine. There was no detectable IEC Axin1 staining in either the colon or small intestine (Fig 2D&2E) of Axin1^ΔIEC^ mice. In addition, we indirectly determined mucus layer thickness and commensal microbiota location by fluorescent in-situ hybridization (FISH). We saw an increase in commensal bacterial invasion into the ileum (Fig 2F) and colon (Fig 2G) of Axin1^ΔIEC^ mice. This data confirms the IEC-specific knockdown of Axin1 and that it contributes explicity to decreased mucus layer thickeness. This Axin1^ΔIEC^ model will allow us to determine the mechanisms in which intestinal epithelial Axin1 regulates intestinal homeostasis and colitis.

**Figure 2.**
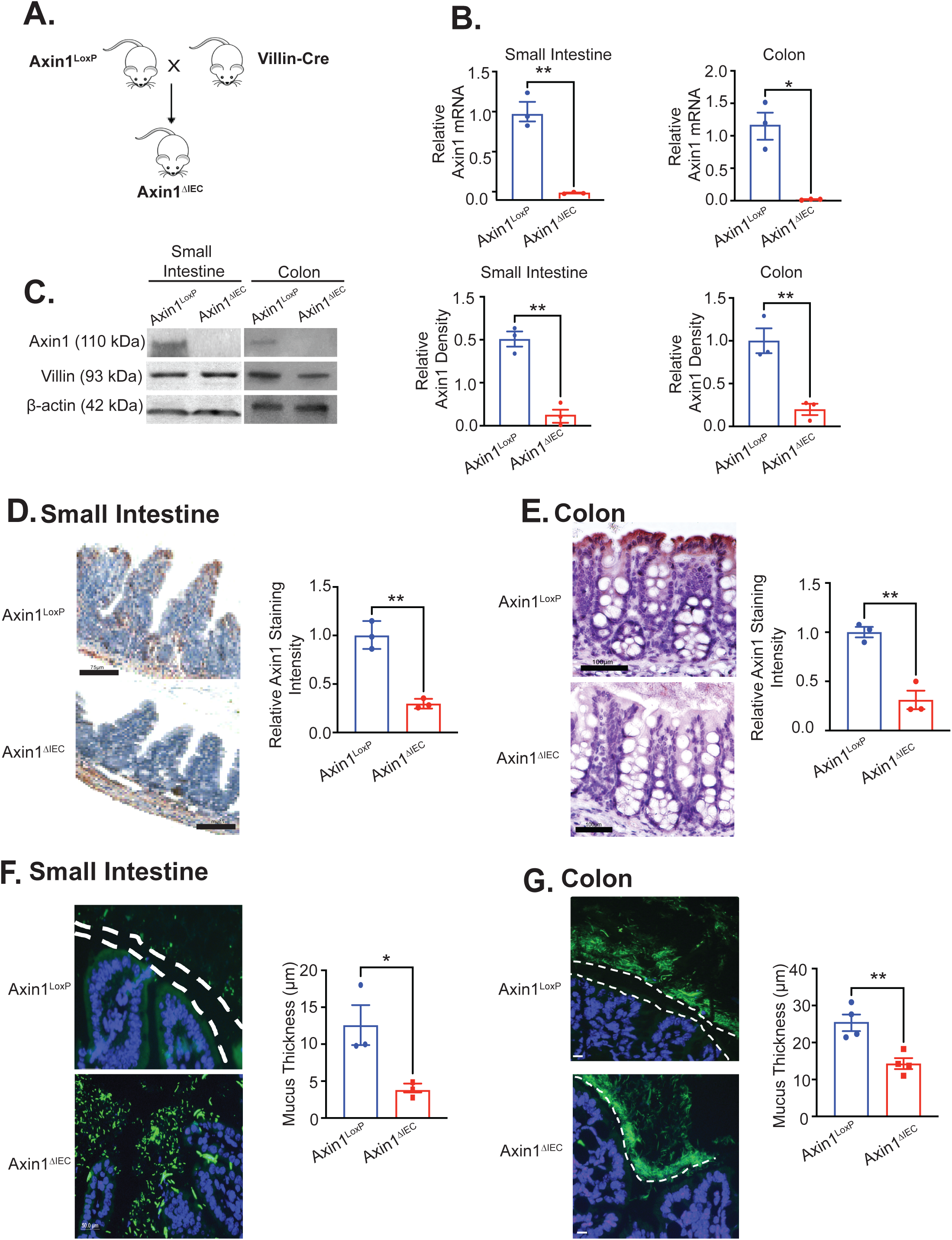
Establishment of Axin1^ΔIEC^ mouse model. (A) Axin1^ΔIEC^ mice, which have Axin1 conditionally knocked out of intestinal epithelial cells were generated by crossing Axin1 ^LoxP /LoxP^ (Axin1^LoxP^) with Villin-cre expressing mice. Villin is expressed only in intestinal epithelial cells. Real-time qPCR of Axin1 mRNA expression in small intestine and colon. (C) Axin1 western blot analysis of intestinal Axin1 with villin as a marker for intestinal epithelial cells. Immunohistochemistry staining of Axin1 in colon (D) and ileum (E) of Axin1^LoxP^ and Axin1^ΔIEC^ mice. Scale bar: 100 µm and 75μm respectively. FISH staining and indirect quantification of mucus thickness of small intestine (F) (n =3) and colon (G) (n = 4) using general bacterial probe EUB388 (green), white dotted lines show indirect thickness of mucus barrier, small intestine scale bar 50μm, colon scale bar 20μm. All aforementioned data is expressed as mean +/-SEM, n=3 per group, Welch’s two-sample t-test used for colon in (B), all other figures were analyzed using unpaired student’s t-test, *P<0.05 and **P<0.01.

### Intestinal Epithelial Axin1 Regulates Intestinal Cell Differentiation and Microbiome

We next determined whether Axin1 deficiency impaired the cell differentiation in the intestine. Axin1^ΔIEC^ mice show a significant increase in Alcian Blue positive goblet cells (GC) (Fig 3A). In addition, there was increased mRNA expression of GC marker, MUC2 in the small intestine of Axin1^ΔIEC^ mice (Fig 3B), suggesting that Axin1 contributes to intestinal GC differentiation. We analyzed the structure of GCs via transmission electron microscopy (TEM) (Fig 3C). Axin1^ΔIEC^ mice had larger mucin granules than Axin1^LoxP^ mice, despite having similar numbers of mucin granules per GC (Fig 3D).

**Figure 3:**
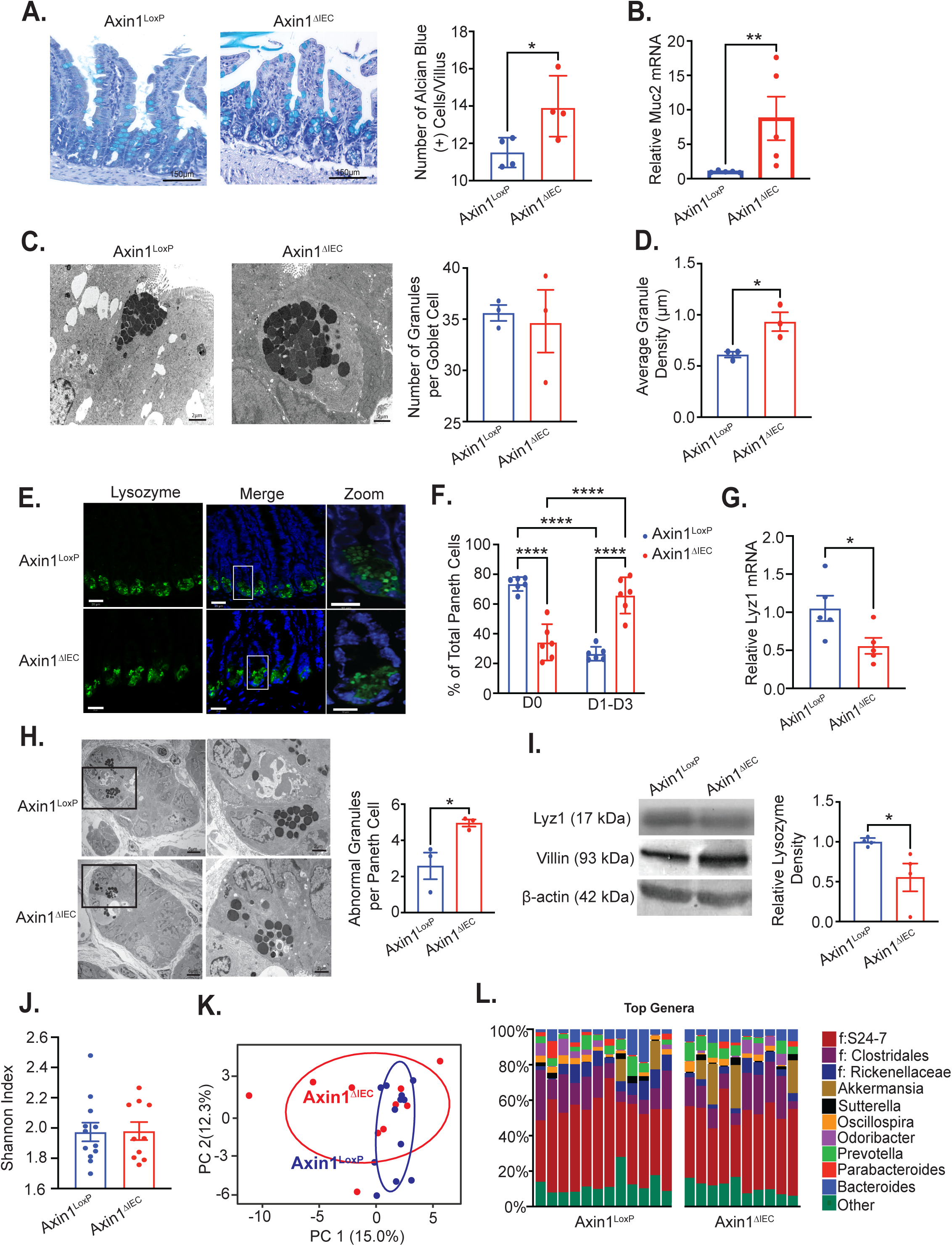
Intestinal Epithelial Axin1 Regulates Goblet Cell Distribution and Paneth cell morphology in the Small Intestine. A) Alcian blue staining of goblet cells in villus of small intestine. Scale bar 150μm (n = 4, >15 villus/mouse). (B) Real-time qPCR mRNA expression of Muc2 in small intestine (n = 5). (C) Axin1^LoxP^ and Axin1^ΔIEC^ goblet cell granules by transmission electron microscopy (TEM). Scale bar 2μm. Goblet cells were identified by presence of cytoplasmic granules and their apical location (n = 3). (D) Average size of mucin granules in Axin1^LoxP^ and Axin1^ΔIEC^ mice (n =3, >15 cells/mouse). (E) Immunofluorescence staining of lysozyme in crypts of small intestine of Axin1^LoxP^ and Axin1^ΔIEC^ mice. Scale bar 20μm, 10μm for zoom image. (F) Percentage of Paneth cells displaying normal (D0) and abnormal (D1-D3) lysozyme morphology (n = 6, >15 crypts/mouse). (G) Lysozyme mRNA in Axin1 mice (n = 6). (H) Lysozyme protein expression by western blot in Axin1 mice (n = 4). (I) Abnormal Paneth cell granules in Axin1 by TEM. Paneth cells were identified by presence of cytoplasmic granules and basal location. Scale bar 6μm, 2μm for zoom image (n = 3, >50 granules/mouse). (J) Shannon diversity index of fecal bacteria between Axin1^LoxP^ and Axin1^ΔIEC^ mice. Data is expressed as mean +/-SEM, n = 10-12 per group, Welch’s t-test. (K) PCA plot visualizing the difference in fecal bacteria between Axin1^LoxP^ (light blue) and Axin1^ΔIEC^ mice (red). The axes explain 27.3% of variation of the separation between the two groups, n = 10-12 per group. (L) Percent abundance of top genera in feces of Axin1^LoxP^ and Axin1^ΔIEC^ mice. All unidentified or other identified species are grouped in “other” as indicated. Genera is color coordinated as indicated by legend. Multiple bars indicate 1 mouse per bar. All aforementioned data is expressed as +/-SEM, two-way ANOVA with Tukey’s method for adjusting multiple comparisons was used in (F) and Wilcoxon rank sumtest was used in (B) and (H), all the analyses in other figures were analyzed using unpaired student’s t-test. *P<0.05, **P<0.01 and ****P<0.0001.

Paneth cells (PCs) are specialized small intestinal epithelial cells that secrete antimicrobial peptides and are crucial for shaping the microbiota and regulating innate immunity [14] [15, 28]). Considering their physiological role, we next assessed PC status in Axin1^ΔIEC^ mice. We categorized PCs based on their lysozyme morphology via immunofluorescent staining described previously by Wu et al[17] (Fig 3E). Normal PCs are indicated as D0, while abnormal PCs are grouped as D1 (disordered), D2 (depleted), and D3 (diffused) lysozyme granule morphologies. We found fewer normal PCs in the Axin1^ΔIEC^ ileum compared to Axin1^LoxP^. Most importantly, we saw a significant increase in the number of abnormal PCs in the Axin1^ΔIEC^ mice (Fig 3F). These abnormal PCs were associated with decreased Paneth Cell (Lyz1) lysozyme at the mRNA level (Fig 3G). We further assessed PC structural morphology by TEM. We saw a significant increase in the number of PCs with less electron-dense granules in the Axin1^ΔIEC^ mice (Fig 3H). Furthermore, decreased lysozyme expression was seen at the protein level (Fig 3I).

### Altered Intestinal Microbiome in Axin1^ΔIEC^ Mice

PCs and GCs are known to modulate the intestinal microbiota profile and function. We examined the bacterial abundance of Axin1^LoxP^ and Axin1^ΔIEC^ mice by 16s rRNA sequencing. We found that the fecal amplicon profile showed unchanged α- and β-diversity between Axin1^LoxP^ and Axin1^ΔIEC^ microbiota (Fig 3J-3K). However, the top genera abundances showed an enrichment in *Akkermansia* and a depletion in Rikenellaceae and *Clostridales* in Axin1^ΔIEC^ mice (Fig 4L). *Akkermansia* is a potential probiotic and a mucolytic bacterium residing in the intestinal mucus layer [29]. These data show that intestinal epithelial Axin1 regulates intestinal secretory cell homeostasis and microbial composition, contributing explicitly to increased abundance of *Akkermansia*.

**Figure 4:**
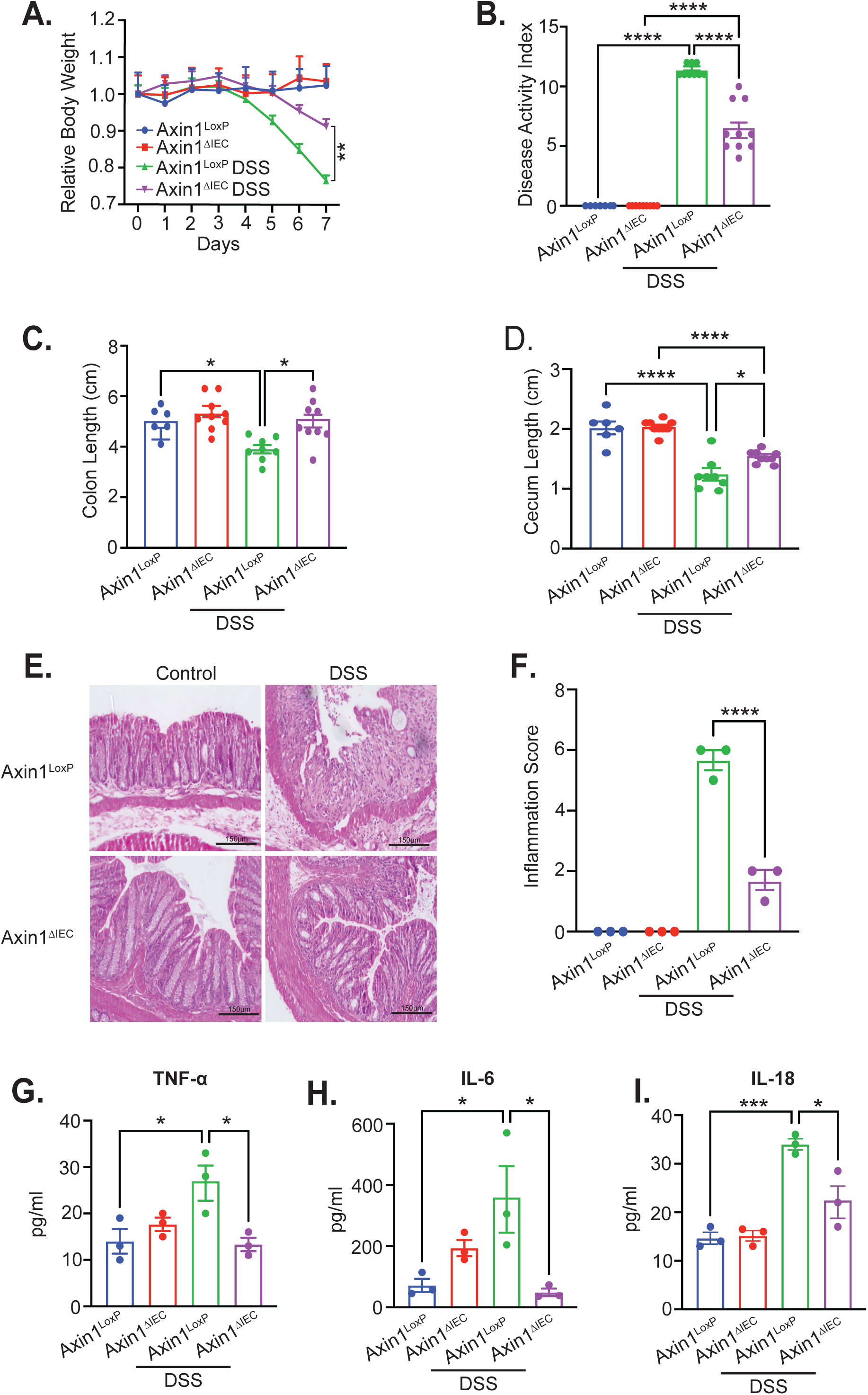
Loss of Intestinal Epithelial Axin1 Confers Protection in DSS-induced Colitis. (A)Relative body weight changes in mice administered 5% DSS for 7 days. Each dot represents minimum of 6 mice. Data expressed as mean +/-SEM, n =6-9, two-way ANOVA, **P<0.01. (B) Colitis severity, (C) colon length and (D) cecum length at day 7 in Axin1 mice. (E) H&E histology of distal colons at day 7. Scale bar 150μm. (F) Histology score of control and DSS-treated mice (n =3). Serum cytokine levels of (G) TNF-α, (H) IL-6 and (I) IL-18 in Axin1 mice (n =3) Aforementioned data is expressed as +/-SEM, two-way ANOVA, *P<0.05, ***P<0.001 and ****P<0.0001.

### Intestinal Epithelial Axin1 Deficient Mice are Less Susceptible to DSS-Induced Colitis

To evaluate the physiological impact of loss of IEC Axin1, we utilized a dextran sodium sulfate (DSS) induced-colitis murine model. Axin1^LoxP^ and Axin1^ΔIEC^ mice were treated with 5% DSS for 7 days, and parameters of colitis were determined. We found Axin1^LoxP^ mice loss more weight during DSS administration compared to Axin1^ΔIEC^ (Fig 4A). Axin1^LoxP^ DSS mice had apparent blood and diarrhea in their stool according to the Disease Activity Index [30] (Fig 4B). Accordingly, colon and cecum shortening was observed in DSS-treated Axin1^LoxP^ mice, compared to DSS-treated Axin1^ΔIEC^ mice (Fig 4C-4D). Colonic histological analysis in DSS-treated mice revealed severe inflammation marked by intestinal epithelial destruction and inflammatory cell infiltration (Fig 4E). In contrast, the histological scores were significantly improved in DSS-fed Axin1^ΔIEC^ mice (Fig 4F). We also tested the serum cytokine profile and specifically found a decrease in TNF-α, IL-6, and IL-18 in Axin1^ΔIEC^ mice compared to DSS-treated Axin1^LoxP^ mice (Fig 4G-I). These data strongly suggest a reduction in inflammation in mice lacking IEC Axin1.

### Cohousing Axin1^ΔIEC^ Mice with Axin1^LoxP^ Mice Increased Susceptibility to DSS Colitis

Axin1^ΔIEC^ mice exhibit alterations in their gut microbiota associated with improved intestinal inflammation outcomes. We hypothesized genetic deletion of Axin1 altered the microbiome contributing to their non-colitogenic phenotype. We co-housed Axin1^ΔIEC^ and Axin1^LoxP^ mice for 4 weeks, followed by 5% DSS challenge for 7 days. We found that co-housing decreased the body weight and DAI of co-housed DSS Axin1^ΔIEC^ mice (Fig 5A-5B). Co-housing also shortened the colon and cecum in Axin1^ΔIEC^ mice to levels like the Axin1^LoxP^ mice (Fig 5C-5D). *Akkermansia muciniphila* in Axin1^ΔIEC^ mice were decreased after co-housing (Fig 5E). Co-housed Axin1^ΔIEC^ mice showed increased intestinal epithelial damage than Axin1^ΔIEC^ mice housed alone (Fig 5F-5G). DSS-treated Axin1^ΔIEC^ mice had decreased serum TNF-α, IL-6, and IL-18. However, co-housing increased serum cytokine levels as seen in the Axin1^LoxP^ mice (Fig 5H-5J). Our results indicate the resistance of Axin1^ΔIEC^ mice to DSS-induced colitis depends on their gut microbiota.

**Figure 5:**
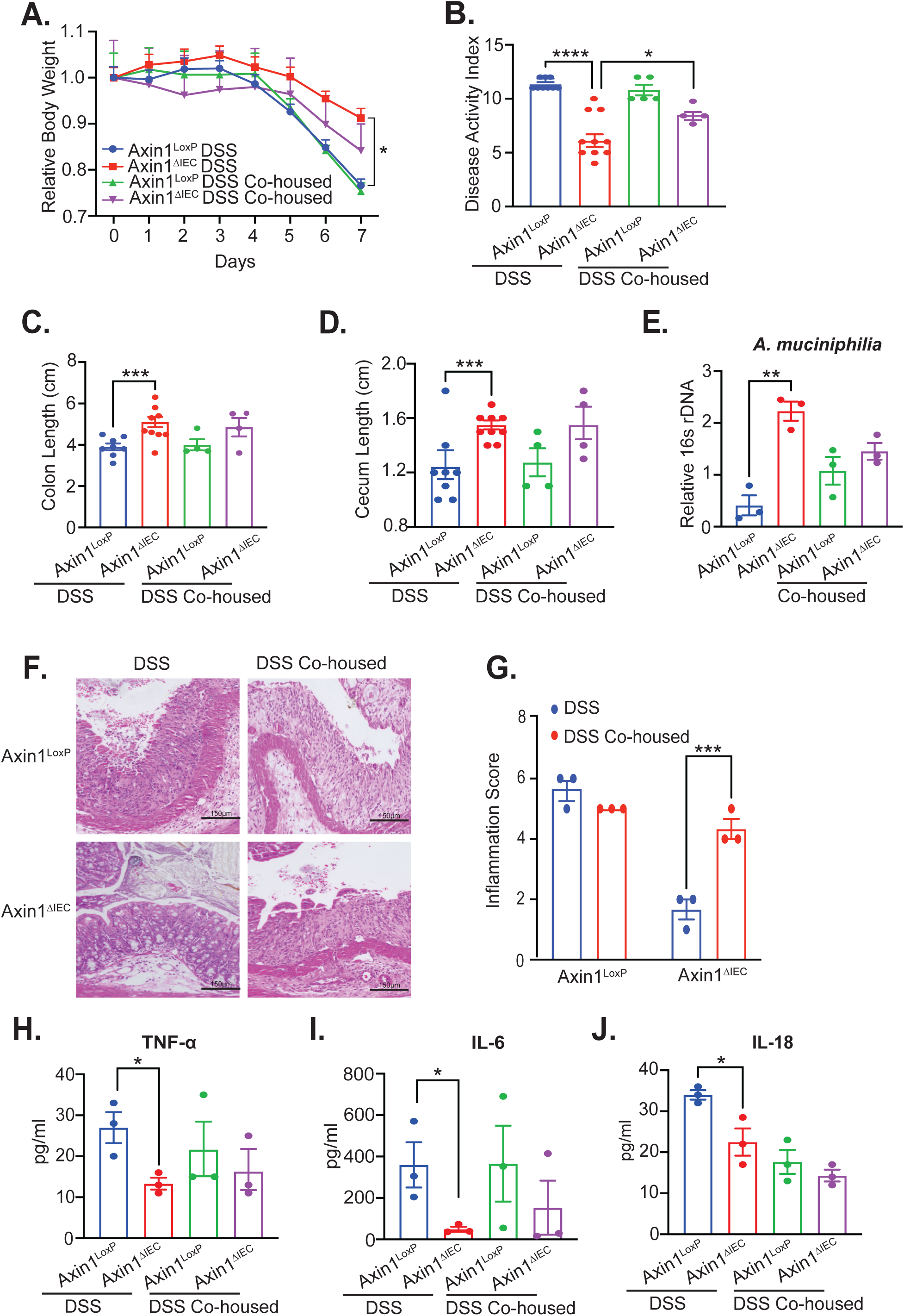
Increased Susceptibility to DSS-induced Colitis in Axin1^ΔIEC^ mice after cohousing with Axin1^LoxP^ mice. Axin1^LoxP^ and Axin1^ΔIEC^ mice were co-housed for 4 weeks and then treated with 5% DSS for 7 days. (A) Relative body weight changes in mice with DSS. Each dot represents minimum of 4 mice. Data expressed as mean +/-SEM, n =4-8, two-way ANOVA, *P<0.05 (B) Colitis severity, C) colon length and (D) cecum length at day 7 in Axin1 mice. Aforementioned data expressed as +/-SEM, n =4-8, two-way ANOVA, *P<0.05, ***P<0.001 and ****P<0.0001. (E) qRT-PCR of *Akkermansia muciniphilia* rDNA expression in Axin1 mice housed alone and co-housed, n =3, +/-SEM, one-way ANOVA, **P<0.01. (F) H&E histology of distal colons at day 7. Scale bar 150μm.(G) Histology score of Axin1 DSS mice, n =3, +/-SEM, One-way ANOVA. ***P<0.001. (H) TNF-α, (I) IL-6 and (J) IL-18 serum cytokines in DSS Axin1^LoxP^ and DSS Axin1^ΔIEC^ DSS mice n =3, +/-SEM, unpaired student’s t test compared between Axin1^LoxP^ and Axin1^ΔPC^ mice, *P<0.05.

### Paneth cell Specific Deletion of Axin1 Resulted in Alteration of Intestinal Secretory Cell Lineages

PCs are known to influence the microbiome’s composition [31, 32] and play a critical role in the intestinal homeostasis and inflammation [14]. We examined the impact of Axin1 PC on intestinal differentiation by generating an Axin1^ΔPC^ mouse model (Fig 6A). We next isolated PCs by labeling them with an anti-CD24 antibody and sorting them into CD24+Paneth cells and CD24-non-Paneth cell fractions (Fig 6B). We tested Axin1 mRNA expression in the isolated Paneth cells by qRT-PCR. PC Axin1 and lysozyme expression in Axin1^ΔPC^ mice was reduced compared to Axin1^LoxP^ PCs (Fig 6C-6D). Immunofluorescent staining showed PCs from Axin1^ΔPC^ mice have abnormal lysozyme morphology compared to Axin1^LoxP^ (Fig 6E-6F). Alcian blue staining showed that Axin1^ΔPC^ mice had increased GCs in the small intestine (Fig 6G). Our data indicate that loss of PC Axin1 results in altered secretory cell differentiation like what we observed in the Axin1^ΔIEC^ model.

**Figure 6:**
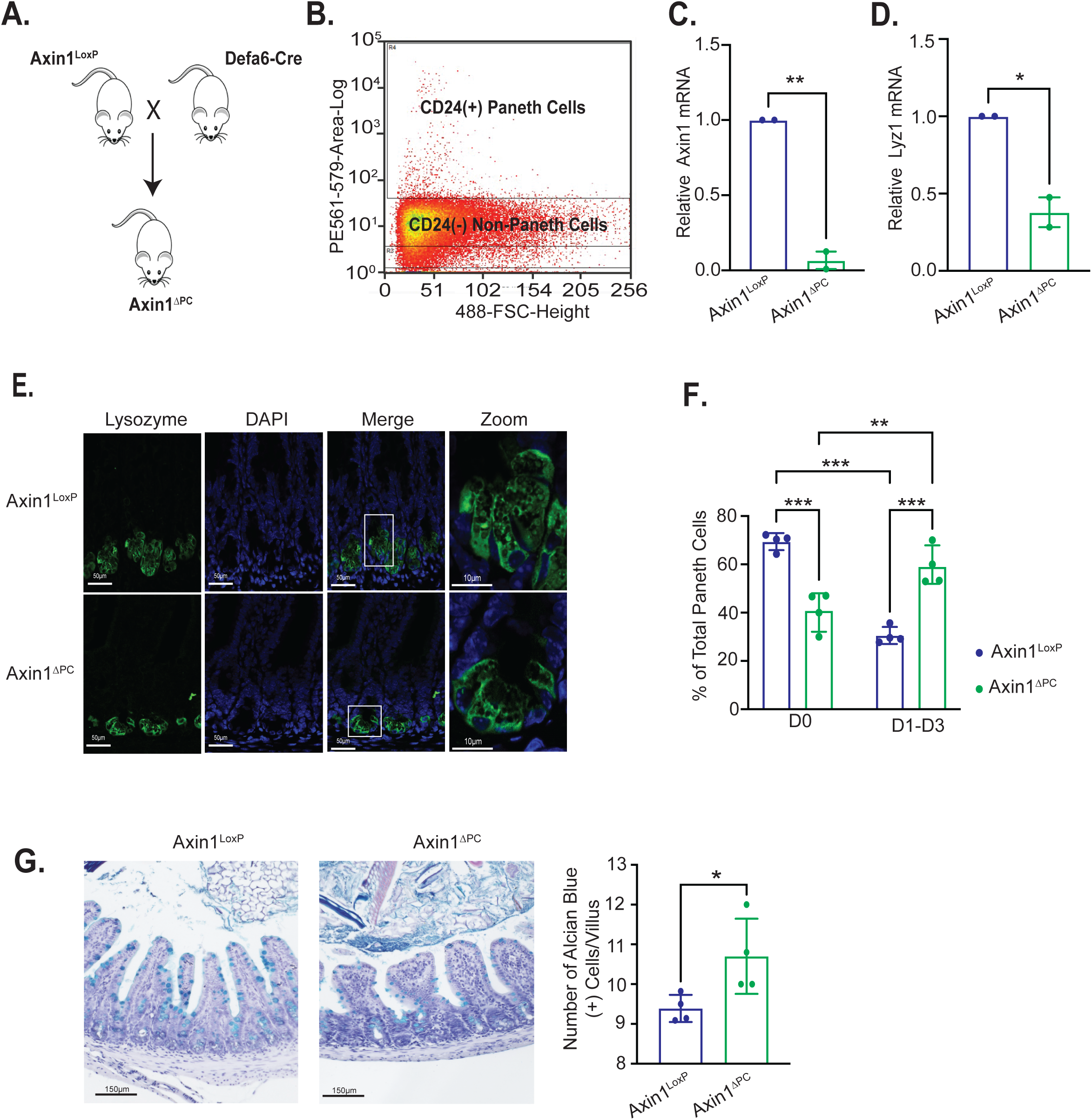
Generation of Axin1^ΔPC^ mice with altered Paneth cell morphology and Goblet cell distribution. (A) Crossing Axin1^LoxP^ mice with DEFA6-cre mice generates a Paneth cell specific knockout of Axin1 (Axin1^ΔPC^). Defensin 6 is only expressed in Paneth cells. (B) Small intestinal epithelial cells were digested into single cells, then stained with anti-CD24 ab and sorted by flow cytometry, n = pool of 3 mice (C) Paneth cell Axin1 knockdown was confirmed by qRT-PCR from CD24+ cells, n = pool of 3 mice (D) Lysozyme mRNA expression of Axin1^LoxP^ and Axin1^ΔIEC^ isolated Paneth cells, n = pool of 3 mice (E) Immunofluorescence staining of lysozyme in crypts of small intestine of Axin1^LoxP^ and Axin1^ΔPC^ mice. Scale bar 50μm, 10 μm for zoom image. (F) Increased number of abnormal (D1-D3) lysozyme morphology in Paneth cells of Axin1^ΔPC^ mice, n =4, >15 crypts/mouse. (G) Alcian blue staining of goblet cells in villus of small intestine in Axin1^LoxP^ and Axin1^ΔPC^ mice, scale bar 150μM, n =4. All aforementioned data is expressed as +/-SEM, two-way ANOVA with Tukey’s method for adjusting multiple comparisons was used to analyze the two factors among four groups in (F) and unpaired student’s t-test was used to compare the differences between two groups in all other figures. *P<0.05, **P<0.01 and ***P<0.001.

### Axin1^ΔPC^ Mice are Less Susceptible to DSS-Induced Colitis and are Dependent on the Gut Microbiota

We next determined the impact of loss of PC Axin1 in the disease progression of DSS-colitis. Axin1^ΔPC^ mice were less susceptible to weight loss and severe disease after DSS administration (Fig 7A-7B). Axin1^ΔPC^ mice have longer colons than Axin1^LoxP^ mice (Fig 7C). Like Axin1^ΔIEC^ DSS mice, Axin1^ΔPC^ DSS mice have improved histologically inflammation and decreased IL-6 (Fig 7D-7F). Because the microbiota was shown to be a key player in Axin1^LoxP^ mice’s vulnerability to DSS-colitis, we examined the transmissibility of the phenotype by performing a co-housing experiment. We challenged Axin1^LoxP^ and Axin1^ΔPC^ mice separately housed and co-housed with DSS. After co-housing Axin1^ΔPC^ with Axin1^LoxP^mice, Axin1^ΔPC^ mice become more susceptible to DSS injury (Fig 7G). Co-housing increased the DAI and shortened the colons of Axin1^ΔPC^ mice to levels like those seen in the Axin1^LoxP^ mice (Fig 7H-7I). Furthermore, we found an increase in fecal *Akkermansia muciniphilia* in the Axin1^ΔPC^ mice, which was significantly reduced after co-housing (Fig 7J). Like Axin1^ΔIEC^ mice, Axin1^ΔPC^ mice had worse histological inflammation and increased levels of IL-6 after co-housing. This data indicates that lack of PC Axin1 may be a driving factor in protection against DSS-induced colitis.

**Figure 7:**
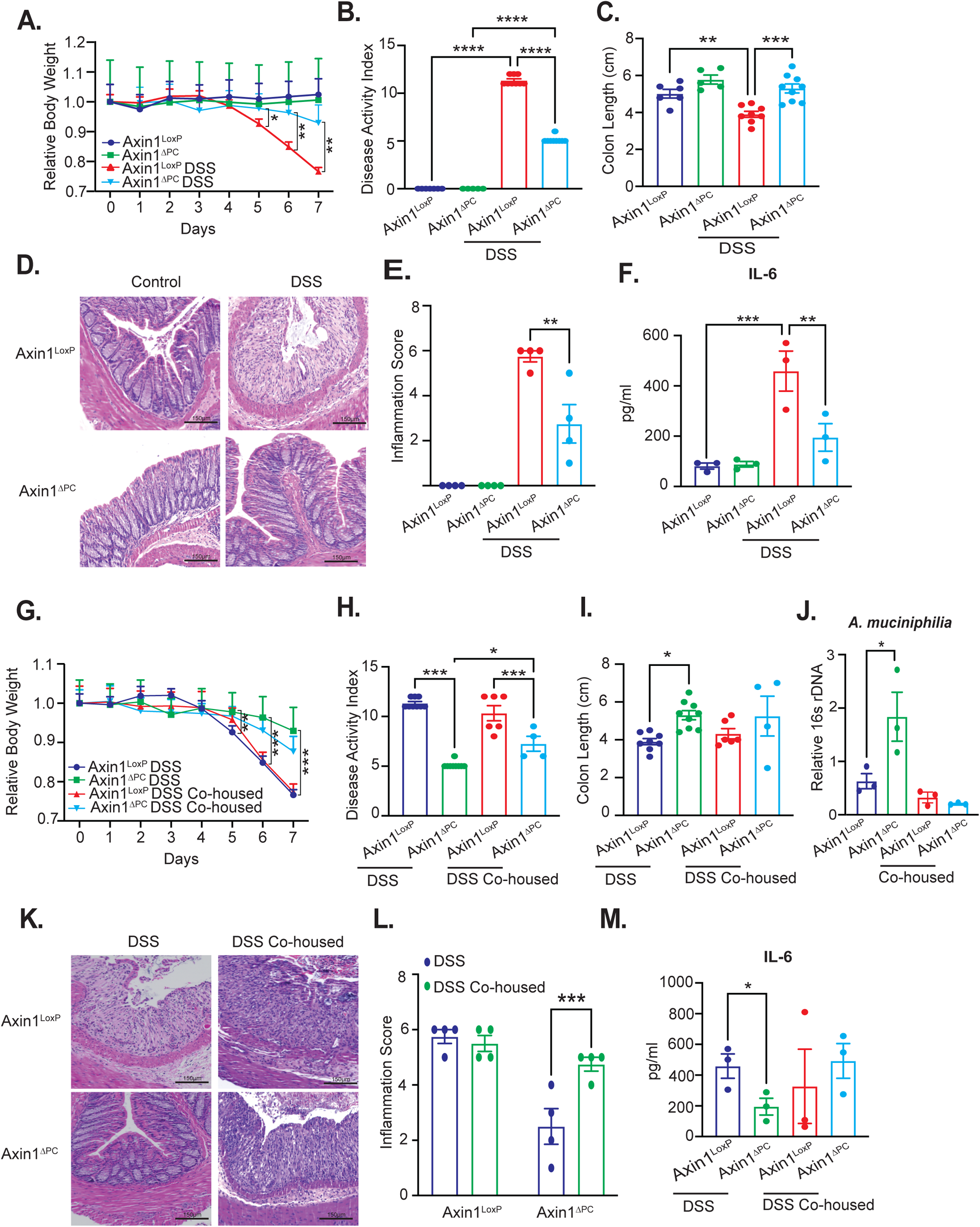
Axin1^ΔPC^ mice are resistant to DSS-induced Colitis and Susceptible after Co-housing with Axin1^LoxP^ mice. Axin1^LoxP^ and Axin1^ΔPC^ mice were co-housed for 4 weeks and then treated with 5% DSS for 7 days. (A) Relative Body weight change during DSS administration. Axin1^LoxP^ mice. Each dot represents minimum 5 mice, two-way ANOVA, *P<0.05 and **P<0.01 (B) Colitis severity and (C) colon length at day 7 in Axin1 mice, one-way ANOVA, **P<0.01, ***P<0.01, and ****P<0.01. (D) H&E histology of distal colons at day 7, scale bar 150μm. (E) Histology score of distal colons at day 7, n =4, +/-SEM, One-way ANOVA, **P<0.01. (F) IL-6 serum cytokine levels in Axin1 mice. n = 3, data expressed as +/-SEM, two-way ANOVA **P<0.01, ***P<0.001. (G) Axin1^LoxP^ and Axin1^ΔIEC^ mice were co-housed for 4 weeks and then treated with 5% DSS for 7 days. Relative body weight changes in mice with DSS. Each dot represents minimum of 4 mice, two-way ANOVA, **P<0.01 and ***P<0.01 (H) Colitis severity and (I) colon length at day 7. Aforementioned data is expressed as n =4-9, +/-SEM, two-way ANOVA, *P<0.05 and ***P<0.001. (J) qRT-PCR of *Akkermansia muciniphilia* rDNA level in Axin1 mice housed alone and co-housed. n =3, +/-SEM, one-way ANOVA, *P<0.05. (K) H&E histology, scale bar 150μm and (L) histology score at day 7 of DSS administration, n =4, +/-SEM, two-way ANOVA. ***P<0.001 (M) Serum IL-6 levels in Axin1 mice housed alone and co-housed, n =3, +/-SEM, unpaired student’s t-test compared between Axin1^LoxP^ and Axin1^ΔPC^ mice, *P<0.05, **P<0.01.

## Discussion

We have demonstrated that loss of intestinal Axin1 leads to altered intestinal epithelial biology, including an increase in goblet cells and changes in Paneth cell morphology and lysozyme expression. Changes in the intestinal epithelium in Axin1^ΔIEC^ mice are specifically associated with an altered gut microbiota favoring increased diversity of beneficial microbes, such as *Akkermansia muciniphilia*. Increase in *Akkermansia muciniphilia* in Axin1^ΔIEC^ mice was associated with thinning of the mucus barrier. Once induced with DSS, Axin1^ΔIEC^ mice had less inflammation than Axin1 sufficient animals. Interestingly, co-housing of Axin1^LoxP^ and Axin1^ΔIEC^ mice reduced the *Akkermansia muciniphilia* abundance resulting in vulnerability to DSS colitis. Axin1^ΔPC^ mice had similar alterations in small intestinal secretory lineages as seen in the Axin1^ΔIEC^ mice. They had protection against DSS-induced colitis and were more susceptible to colitis after being co-housed with Axin1^LoxP^ mice (Fig 9).

In the current study, we have established two unique experimental models to study the role of Axin1 in intestinal function. Because complete inactivation of Axin1 leads to early embryonic lethality, it is impossible to identify the potential role of Axin1 in later developmental processes using global Axin1 knockout [8]. To investigate the tissue-specific function of Axin1, we developed a system in which Axin1 is conditionally deleted from IECs and PCs utilizing cre-recombinase driven under the villin and defensin6 promoters, respectively. These models allow us to understand the fundamental role of Axin1 in intestinal and microbial homeostasis and in host responses to inflammatory stimulators.

Our data from Paneth cell Axin1 deleted mice have demonstrated the critical role of Axin1 in innate immunity. We have shown that Paneth cell Axin1 confers protection against DSS-induced colitis and the gut microbiota contributes to the non-colitogenic phenotype in Axin1^ΔPC^ mice. PCs play a role in shaping the gut microbiota including secreting antimicrobial peptides, like lysozyme. Increased lysozyme mRNA was observed in IECs of patients with UC. In addition, lysozyme expression is correlated with the degree of intestinal inflammation[33]. It has also been shown that mice lacking PC lysozyme (Lyz1^-/-^) have alterations in their gut bacterial landscape. This alteration was associated with protection from DSS colitis and increased mucolytic bacteria. We have shown intestinal Axin1 deficiency produced similar results as Lyz1^-^

^/-^ mice [32]. Axin1 is classically known as a Wnt/β-catenin signaling regulator. Wnt/β-catenin signaling provokes the differentiation and maturation of PCs, thereby regulating antimicrobial peptide production. Mice lacking the Wnt/β-catenin signaling regulator TCF1 demonstrated a decrease in lysozyme [34]. Axin1 may regulate lysozyme expression in a Wnt/β-catenin dependent fashion. Our study has provided new insights into the molecular mechanism that might contribute to inflammation through intestinal epithelial and PC Axin1.

We have identified increased Axin1 expression at the mRNA and protein level in human UC, consistent with a previous study reporting Axin1 serum levels were elevated in patients with UC [35]. Activation of Wnt/β-catenin signaling is an aspect of human IBD and colitis animal models [6, 36]. It is likely that Axin1 may regulate intestinal inflammation and IBD through negative regulation of Wnt/β-catenin signaling. However, Axin1 is a multi-domain protein and has been shown to interact with many proteins in several signaling pathways such as SAPK/JNK, p53, TGFβ, and Wnt signaling [37, 38, 39, 40]. It is unknown how Axin1 may simultaneously regulate multiple signaling pathways in inflammation and infection.

*Akkermansia muciniphilia*, a potential probiotic and only member of the Akkermansia genus, is abundant in the gastrointestinal tract and plays a critical role in maintaining gut and microbial homeostasis and immunity. Our study has shown the direct modulation of Axin1 status on *Akkermansia muciniphilia* in the intestine. Reductions in *Akkermansia muciniphilia* have been demonstrated in human IBD fecal and mucosal samples[41]. Evidence suggests the protective role of *Akkermansia muciniphila* in diseases such as IBD, type 2 diabetes, and obesity [42, 43, 44]. Furthermore, administration of *Akkermansia muciniphilia* ameliorates DSS-induced colitis[30], metabolic disorders[45], and obesity[46] in mice. Mice colonized with *Akkermansia muciniphilia* develop immune tolerances toward commensal bacteria [47]. We found that Axin1^ΔIEC^ mice are protected from DSS-induced inflammation but have a weakened mucus barrier. It’s been shown that a depleted mucus barrier results in susceptibility to DSS-induced colitis [48, 49, 50]. However, increased Muc2 and *Akkermansia muciniphilia* have been shown to protect against DSS-colitis [30, 50], which collectively may allow the Axin1^ΔIEC^ mice to have increased colitis-protection compared to controls. The exact mechanism in which Axin1 regulates the gut microbiota is still unknown. In the future, we will investigate the Axin1-*Akkermansia muciniphilia* axis and a therapeutic role of *Akkermansia muciniphilia* in Axin1 sufficient mice through modulating intestinal mucosal immunity.

In conclusion, our study has demonstrated a novel and critical role of Axin1 in regulating intestinal epithelial development. Paneth cell Axin1 maintains gut and microbial homeostasis which may be the driving factor to protect against colitis. Loss of intestinal Axin1 may alter innate intestinal immunity through its regulation of PC function, which in turn modulates an anti-colitogenic microbiota. Our findings will provide insights into the development of IBD and potential therapeutic strategies for human IBD.

## Acknowledgements/Funds

We would like to acknowledge the VA Merit Award 1 I01BX004824-01, the NIDDK/National Institutes of Health grant R01 DK105118, and R01DK114126 to Jun Sun. R01DK114126 supplement is to promote Diversity in Health-Related Research for PHD student Shari Garrett. We would like to thank Dr. Meena Rao for reading this manuscript and providing insightful suggestions. Figen Seller at the University of Illinois Chicago Electron Microscopy Core for assistance in obtaining transmission electron microscopy images and the University of Illinois Chicago DNAS facility for assistance with DNA sequencing. Working model was created using BioRender. The study sponsors play no role in the study design, data collection, analysis, and interpretation of data. The contents do not represent the views of the United States Department of Veterans Affairs or the United States Government.

## Abbreviations used in this paper

CD: Crohn’s Disease
DSS: Dextran Sulfate Sodium
GI: Gastrointestinal
H&E: Hematoxylin and Eosin
IBD: Inflammatory Bowel Disease
IEC: Intestinal Epithelial Cells
IF: Immunofluorescence
IHC: Immunohistochemistry
LPS: Lipopolysaccharide
LoxP: Locus of X-over, P1
MMP7: Matrix Metalloproteinase 7
MUC2: Mucin 2
OTUs: operational taxonomic units
PC: Paneth Cell
PCA: Principal Coordinate Analysis
qRT-PCR: Quantitative Real Time Polymerase Chain Reaction
UC: Ulcerative Colitis

---

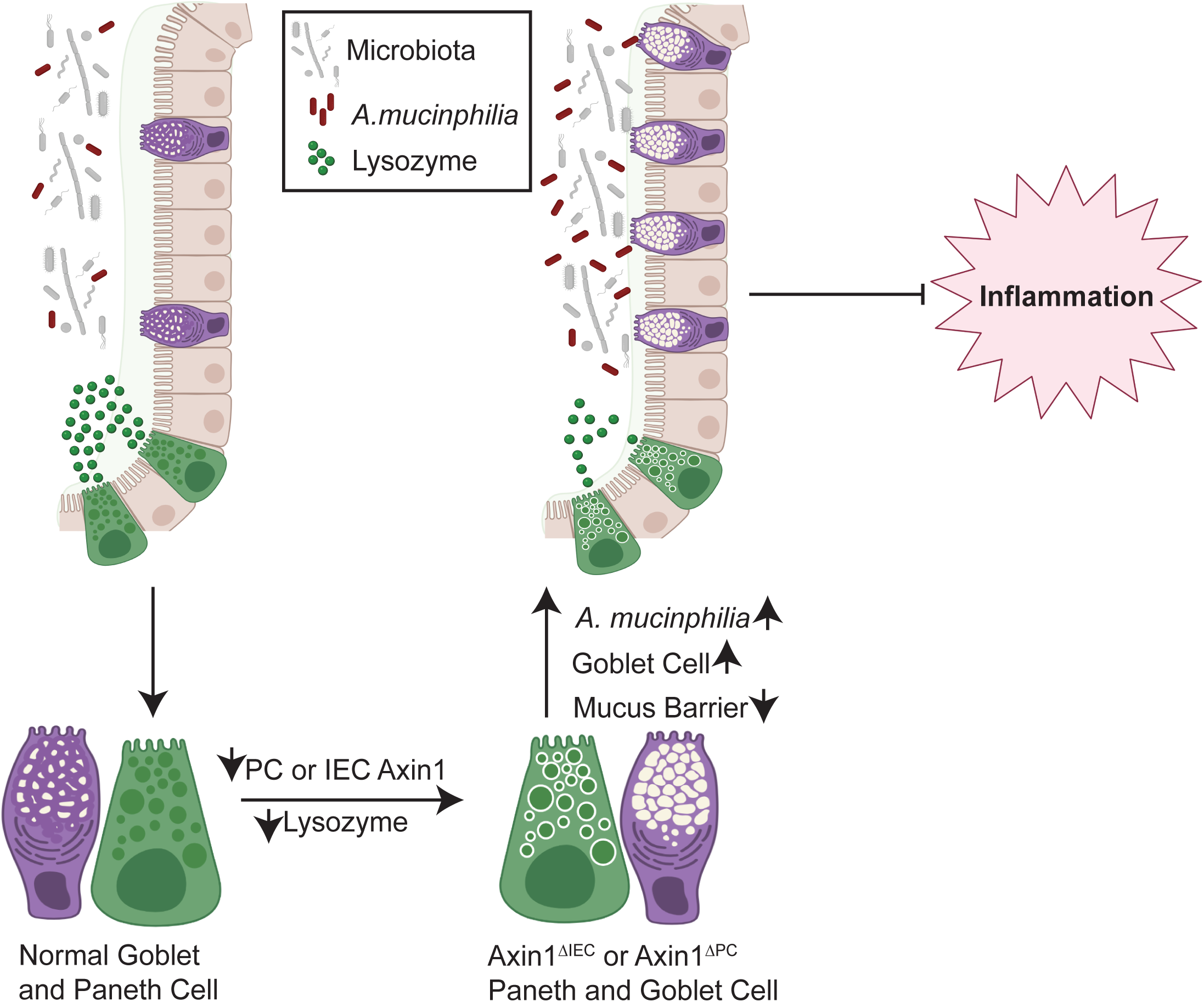

